# Evidence that ultrafast non-quantal transmission underlies short-latency vestibular evoked potentials

**DOI:** 10.1101/2023.05.15.540767

**Authors:** Christopher J. Pastras, Ian S. Curthoys, Mohsen Asadnia, David McAlpine, Richard D. Rabbitt, Daniel J. Brown

**Affiliations:** Faculty of Science and Engineering, School of Engineering, Macquarie University, Sydney, NSW, 2109, Australia; Vestibular Research Laboratory, The University of Sydney, School of Psychology, Sydney, NSW, 2050, Australia; Department of Linguistics, The Australian Hearing Hub, Macquarie University, Sydney, NSW, 2109, Australia; Departments of Biomedical Engineering, Otolaryngology, and Neuroscience Program, University of Utah, Salt Lake City, UT, 84112, USA; School of Pharmacy and Biomedical Sciences, Curtin University, Bentley, Western Australia 6102, Australia

**Keywords:** non-quantal, ephaptic, synaptic transmission, inner ear, vestibular, cochlea, forward masking, latency

## Abstract

Amniotes evolved a unique calyceal postsynaptic terminal in the vestibular organs of the inner ear that underpins quantal and non-quantal transmission at the synapse of sensory hair cells and vestibular afferent neurons. The non-quantal component is of particular interest as it includes an ultrafast synaptic current thought to underlie the exquisite synchronization of action potentials in vestibular afferent fibres to dynamic stimuli such as sound and vibration. Here we demonstrate evidence that non-quantal transmission is responsible for short latency vestibular evoked potentials (vCAPs) in the guinea pig utricle. We first show that, unlike auditory nerve responses which are completely abolished, vCAPs are insensitive to local administration of the AMPA receptor agonist CNQX. Moreover, latency comparisons between presynaptic hair cell and postsynaptic neural responses reveal that the vCAP occurs without measurable synaptic delay. Finally, using a paired-pulse stimulus designed to deplete the readily releasable pool of synaptic vesicles in hair cells, we reveal that forward masking is lacking in vestibular responses, compared to the equivalent cochlear responses. Our data support the hypothesis that the fast component of non-quantal transmission at calyceal synapses is indefatigable and responsible for ultrafast responses of vestibular organs evoked by transient stimulation.

**Significance:** The mammalian vestibular system drives some of the fastest reflex pathways in the nervous system, ensuring stable gaze and postural control for locomotion on land. To achieve this, terrestrial amniotes evolved a large, unique calyx afferent terminal which completely envelopes one or more pre-synaptic vestibular hair cells, which transmits mechanosensory signals mediated by quantal and nonquantal (NQ) synaptic transmission. We present several lines of data in the guinea pig that reveal the pre-synaptic transmission of the most sensitive vestibular afferents are faster than their auditory nerve counterparts. Here, we present neurophysiological and pharmacological evidence that this vestibular speed advantage arises from ultrafast NQ electrical synaptic transmission from Type I hair cells to their calyx partners.

## Introduction

The vestibular system evolved over 400 million years ago in primitive fish (Higuchi et al., 2019), and is the evolutionary precursor to the modern mammalian cochlea (Manley, 2012). The five peripheral vestibular sensory organs detect linear and angular accelerations in three-dimensional space (Curthoys et al., 2017; Goldberg et al., 2012). Fibres from all sense organs travel to the brainstem and cerebellum and terminate in the vestibular nuclei. In these secondary neurons, the vestibular system interfaces with sensory and motor systems, contributing to several levels of nervous function, including some of the fastest reflexes in biology designed to maintain visual and postural stability during dynamic head and body movements.

Collectively, primary vestibular afferent neurons provide the central nervous system with information spanning a broad frequency bandwidth and dynamic range, with individual neurons specialized to encode specific directional and temporal features of diverse gravito-inertial stimuli. Afferent fibres innervating the utricle, precisely encode the timing of transient inertial stimuli, and generate precisely timed action potentials to sinusoidal stimulation (Curthoys et al., 2016; McCue and Guinan, 1994) up to ∼3kHz with a temporal fidelity exceeding that of auditory spiral ganglion neurons in the same species (Curthoys et al., 2019; Palmer and Russell, 1986). These sensitive vestibular neurons make calyceal synaptic contacts on Type I hair cells in the striolar region of the sensory epithelium (Curthoys et al., 2006; Goldberg *et al*., 2012; Lysakowski et al., 2011). The postsynaptic calyceal terminal responsible for this high-fidelity transmission is unique to amniote vestibular organs and supports three distinct forms of neurotransmission: i) glutamatergic quantal transmission analogous to that responsible for synaptic transmission of sound information between Type I spiral ganglion neurons and cochlear inner hair cells (Highstein et al., 2015; Matsubara et al., 1999; Rennie and Streeter, 2006; Ruel et al., 1999; Rutherford et al., 2021), ii) K^+^ build up in the synaptic cleft responsible for a slow non-quantal (NQ) synaptic current (Contini et al., 2020; Contini et al., 2017; Govindaraju et al., 2021; Govindaraju et al., 2023; Highstein et al., 2014; Holt et al., 2007; Lim et al., 2011), and 3) electrical coupling between the hair cell and the terminal responsible for an ultrafast NQ current (Contini *et al*., 2020; Songer and Eatock, 2013). Here, we examine the hypothesis that the ultrafast electrical NQ current is responsible for short latency responses to transient stimuli in the guinea pig utricle.

Ultrafast electrical transmission between the Type I hair cell and the calyx terminal has been reported for calyceal synapses in turtle posterior semicircular canal based on paired pre- and postsynaptic recordings (Contini *et al*., 2020)—and this is consistent with recordings in immature (P1-4) rat saccular calyceal terminals, suggesting the presence of an ultrafast NQ component (Songer and Eatock, 2013). We hypothesized that the same NQ current is present in adult mammals, and underlies short latency neural responses, including Vestibular stimulus Evoked Potentials (VsEPs) commonly used for vestibular phenotypic screening in small amniotes (Jones and Jones, 1999; Jones and Lee, 2021). VsEPs are dominated by vestibular Compound Action Potentials (vCAP) generated by synchronized firing of many vestibular afferent neurons (Brown et al., 2017; Goldstein and Kiang, 1958; Pastras et al., 2023).

We first confirmed the possibility of fast, non-quantal transmission by perfusing glutamate blockers into the inner ear of our guinea pig model and comparing the effects on vestibular and cochlear synaptic transduction. Second, we examined and compared the latency of vCAPs to cCAPs recorded from the guinea pig. Finally, we examined the effects of a forward masking paradigm, in which paired stimuli are presented in short succession with the interval between the pulses controlled (paired-pulse interval, PPI). Here, we aimed to examine the presence of the classically described phenomenon of forward masking (Goldstein and Kiang, 1958) present in the cochlea, which is thought to arise primarily from the kinetics of depletion/replenishment of the readily releasable pool (RRP) of synaptic vesicles within inner hair cells (Peterson et al., 2014).

## Methods

### Animal preparations and surgery

Experiments were performed on 25 adult tri-coloured guinea pigs (*Cavia porcellus*) of either sex, weighing between 300-500g. All procedures were approved by the University of Sydney Animal Ethics Committee (Protocol# 2019/1533). All animals had a positive Preyer’s reflex indicating good hearing. This was further confirmed via inspection of the middle ear cavity under the surgical microscope, and later via auditory cCAP thresholds. Prior to surgeries, guinea pigs received pre-anaesthetic medications of Atropine sulfate (600μg in 1ml, Pfizer, AUS), and Buprenorphine HCl (Temgesic, 300ug in 1ml, Indivior, New Zealand). Thereafter, guinea pigs were transferred to a Perspex induction chamber and were anaesthetized using 2– 4 % isoflurane (99.9% Isoflurane Liquid Inhalation; Henry Schein, US). Once lacking a foot-withdrawal reflex, animals were transported to the surgical table to be tracheotomized and artificially ventilated with a mixture of oxygen and isoflurane (2%). Heart rate and blood oxygenation (SpO2) levels were continuously monitored (Nonin Medical Inc., MN, USA), with the animal’s core temperature regulated via a heating pad (Kent Scientific, CT, USA). Animals were then mounted between custom-made ear bars and were electrically grounded via a Ag/AgCl electrode placed in the neck musculature.

### Stimuli and recordings

All stimuli and responses were generated and recorded using customized LabVIEW programs (National Instruments, TX, USA). Stimuli were produced using an external soundcard (SoundblasterX7, Creative Inc., Singapore). Physiological responses were amplified by 10,000x (80dB) with a 1-Hz to 10-kHz band-pass filter (IsoDAM, WPI, FL, USA), prior to being digitized at 40 kHz and 16-bit using an Analog-to-Digital Converter (NI9205, National Instruments, TX, USA).

To record neural responses from the vestibular system, the dorsolateral bulla was opened, and a non-inverting Ag/AgCl electrode was inserted into the bony facial nerve canal near the superior branch of the vestibular nerve. The inverting reference electrode was inserted nearby in the neck musculature. Auditory potentials such as cochlear nerve CAP and Cochlear Microphonic were recorded from the round window using air conducted sound (ACS) or bone conducted vibration (BCV) pulses. Linear acceleration transients were delivered to the skull via BCV pulses using a Brüel & Kjær minishaker (Type 4810, Denmark), which was rigidly coupled to the contralateral ear-bar. A 3-axis accelerometer (830M1; flat frequency response >15kHz; TE Connectivity, Switzerland) was secured to the ear-bar and was used to quantify input stimuli in G (1G = 9.81 m/s2).

### Cochlear ablation and utricular exposure

The cochlea was accessed via a ventral surgical approach, as previously described in (Pastras et al., 2017; Pastras et al., 2018a; Pastras et al., 2021; Pastras et al., 2020) (see Fig. 4A). Fine-tip forceps, a hooked-needle, and scalpel were used to surgically ablate the cochlea, starting at the apex, and moving to the base. Tissue wicks were inserted to absorb fluid and blood following ablation. Cochlear tissue and bone were cleared to expose the basal surface of the utricular macula (Fig. 1E, 4A). Tissue wicks were implanted adjacent to the macula to control fluid build-up on the epithelium.

**Figure 1:**
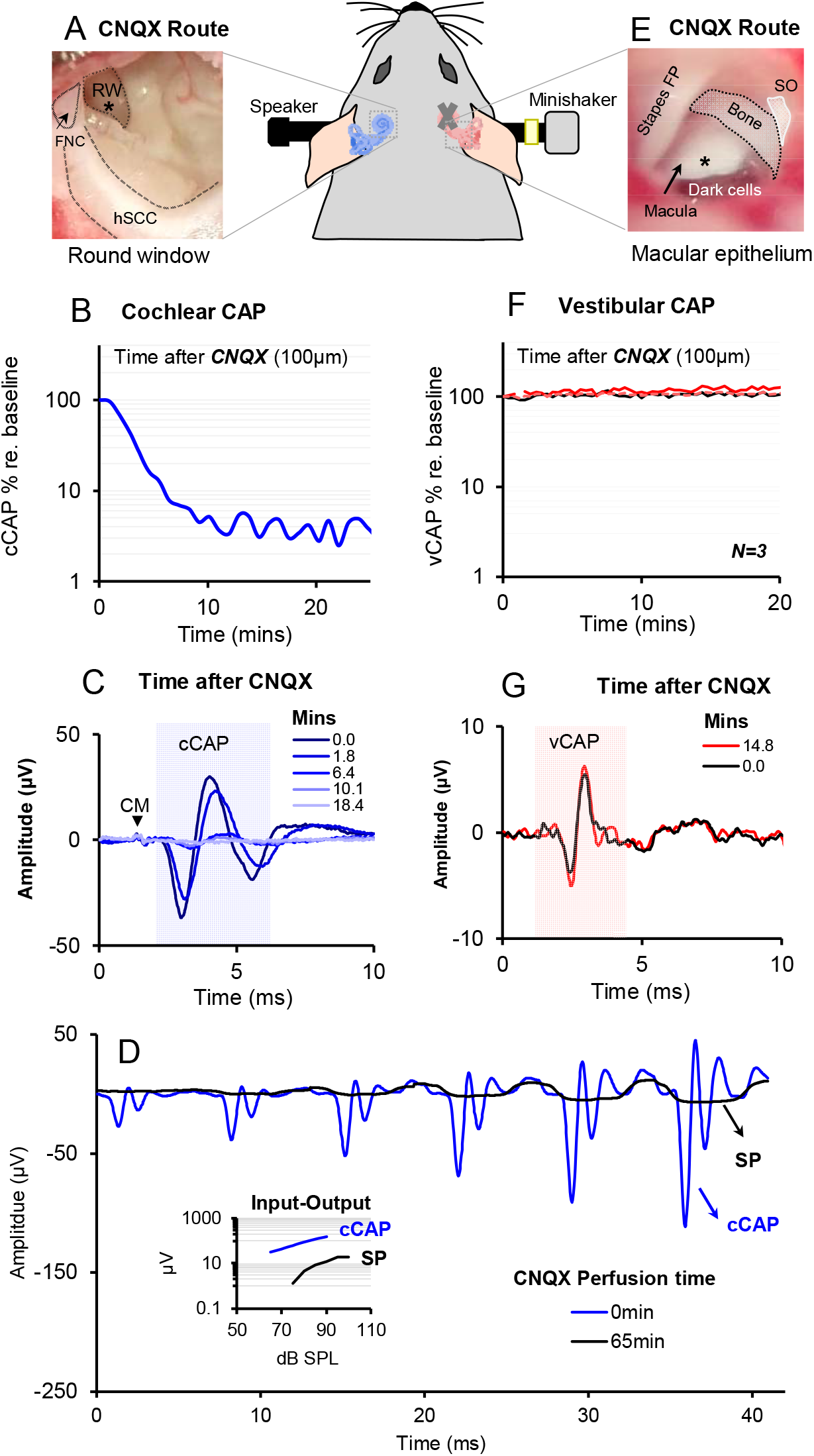
Effect of 6-cyano-7-nitroquinoxaline-2,3-dione (CNQX) on Cochlear vs Vestibular CAPs in the guinea pig. CNQX (100μM in artificial perilymph) was applied to the A) cochlear round window, and d) the utricle via the macular epithelium (approximate drug delivery locations denoted by asterisks). B, F) Time chart of cCAP and vCAP amplitudes normalized as a percentage of baseline. The cCAP was diminished whilst vCAPs remained unperturbed demonstrating differential effects of CNQX on cochlear vs vestibular afferent function. C, G) Averaged cCAP and vCAP responses (100 stimulus presentations) corresponding to labelled time points after CNQX administration. D) CNQX blocks auditory CAPs (blue) but not the presynaptic SP (black). Inset: Input-Output growth functions of auditory CAPs before and SPs after CNQX perfusion. Abbreviations: RW = Round Window, FNC = Facial Nerve Canal, hSCC = horizontal Semicircular Canal, Stapes FP = Stapes Footplate, SO = Saccular Otoconia, CM = Cochlear Microphonic, cCAP = cochlear Compound Action Potential, vCAP = vestibular Compound Action Potential.

### Application of 6-cyano-7-nitroquinoxaline-2,3-dione (CNQX)

A 10mM stock solution of CNQX was first produced by dissolving 50mg of CNQX disodium salt (molecular weight = 276.12, ab120044, abcam, Sydney, AUS) in 18mL of artificial perilymph (NaCl, 130; KCl, 4; MgCl_2_, 1; CaCl_2_, 2; pH 7.3; osmolarity 320 mOsmol). Stock solutions were stored as aliquots in tightly sealed vials at -20°C, as per supplier guidelines. Aliquots were used within 2 weeks of freezing, prior to the recommended useable period of 1 month. Before use, and prior to opening the vial, solutions were allowed to equilibrate to room temperature for approximately 1 hour. Thereafter, stock solutions were diluted to 100μM in fresh artificial perilymph. Approximately 0.1mL of 100μM CNQX/Artificial perilymph solution was drawn up in a 1ml syringe with a 27-gauge Luer lock needle (Hamilton, MA, USA). 2-3 drops of 100μm CNQX were placed onto the round window or above the perilymphatic layer of the utricular macula under the visual guidance of an operating microscope (corresponding to the approximate location denoted by the asterisk symbol in Fig. 1E). CAP responses were monitored before, during and after application of the drug.

### Electrical stimulation of vestibular neural targets

Central stimulation of the peripheral afferents was chosen over peripheral stimulation, to avoid possible complexities in interpretation of the electrically evoked vestibular compound action potential (evCAP). For example, electrical stimulation of the periphery can result in multiple activation sites of the primary afferent neuron, such as its peripheral axon, soma, or along its central axon, which has been documented in the cochlea (Stypulkowski and Van den Honert, 1984). Moreover, electrodes placed in the periphery can evoke *electrophonic* responses that mimic fibre activation, which arise from the sensory hair cells (Moxon, 1971; Van den Honert and Stypulkowski, 1984). Thirdly, central vestibular stimulation evoking antidromic electrically evoked vCAPs have been shown to fully cancel the orthodromic BCV vCAP (Pastras et al., 2018b), which is evidence that the antidromic electrically evoked vCAP arises from the same neurons as the BCV vCAP. This can be explained by the electrically evoked antidromic action potential colliding with, and cancelling, the vibration-evoked orthodromic action potential in the same afferents. Hence, by stimulating the vestibular afferents at their proximal ending in the central vestibular system, we were confident that the evCAP arose from the same neural targets that generate the BCV evoked vCAP.

To electrically stimulate the proximal endings of the primary vestibular afferents at the central vestibular system, surgery was undertaken to expose the floor of the fourth ventricle for placement of bipolar stimulating electrodes. Specifically, a small incision was made between the caudal point of the occipital bone and Lambda. Thereafter, a posterior craniotomy was made, and the dura mater was cut. A section of the flocculus and paraflocculus were aspirated to fully expose the floor of the fourth ventricle. Fluid build-up was controlled with tissue wicks. A pair of platinum bipolar electrodes (1.0MΩ impedance, 400μm diam.) coated with a parylene-C layer (Microprobes, MD, USA) were used for stimulation of vestibular neural targets. Stimulating electrodes were positioned lateral of the floor of the fourth ventricle under the guidance of a surgical microscope using a 3-axis clamp mount micromanipulator (MM-33, ADInstuments, USA). Electrode placement was based on guinea pig stereotaxic map coordinates (Rapisarda and Bacchelli, 1977; Voitenko and Marlinsky, 1993) and guinea pig immunohistochemistry studies (Motts et al., 2008). The bipolar electrodes were positioned at the same lateral-medial plane as the crossed Olivocochlear bundle, at the midline floor of the fourth ventricle, and positioned lateral of the facial nerve genu beneath the sulcus limitans (Pastras *et al*., 2018b). Low threshold current twitches indicated stimulation of the facial nerve genu or abducens nucleus. To locate appropriate vestibular afferent targets, subsequent dorsolateral electrode adjustment/positioning was required. Final positioning of the bipolar electrodes dorsolateral of the midline, was determined by the maximal physiological effect on the vCAP with the lowest shock intensity, analogous to previous work (Pastras *et al*., 2018b). Current shocks (bipolar, 100μs biphasic pulse width) were delivered via an isolated, biphasic current stimulator (Model DS4, Digitimer Ltd., United Kingdom).

### Data availability

The datasets used during the current study, as well as the code for data acquisition and analysis are available from the corresponding author on reasonable request.

### Results

Experiments in adult guinea pigs were undertaken comparing robust functional cochlear and vestibular responses to determine differences in synaptic transmission modes.

### Effect of CNQX on cochlear vs. vestibular CAPs

To assess the existence of a non-glutamatergic form of synaptic transmission in the vestibular system, we applied the glutamate (AMPA) antagonist CNQX (100μM in artificial perilymph) to the utricular macula (Fig. 1E), as well as to the cochlear round window (Fig. 1A). Here, opposite ears of the same animal were utilised (see Fig. 1). Whilst CNQX abolished any detectable cCAP response within 10 minutes of drug application (Fig. 1B-D), vCAPs persisted even after prolonged CNQX perfusion (Fig. 1F, G). Importantly, hair cell responses (microphonic and summating potentials) in both the cochlea and utricle were unaffected by CNQX (Fig. 1D).

### Vestibular vs. Auditory Response Latency

Next, we compared latencies of the vestibular and cochlear evoked potentials (vCAPs and cCAPs), using the onset of hair cells responses (vestibular microphonic; VM, or cochlear microphonic; CM) as a temporal reference point (Fig. 2B and 2C). Both responses were evoked by a 0.6ms, condensation or rarefaction gaussian ACS pulse, with the polarity of the stimulus resulting in significant differences in the latency of the vCAP. Results reveal that both condensation and rarefaction ACS pulses produced a latency of 0.35ms between the vCAP and VM (Fig. 2C), suggesting a delay of 0.35ms to peak action potential voltage in the guinea pig utricle. In contrast, the latency of the cCAP relative to the CM for a condensation or rarefaction ACS pulse was 1.35ms and 1.28ms respectively, which is approximately 1ms longer than the latency of the vCAP. Absolute latencies are compared in Fig. 2D.

**Figure 2:**
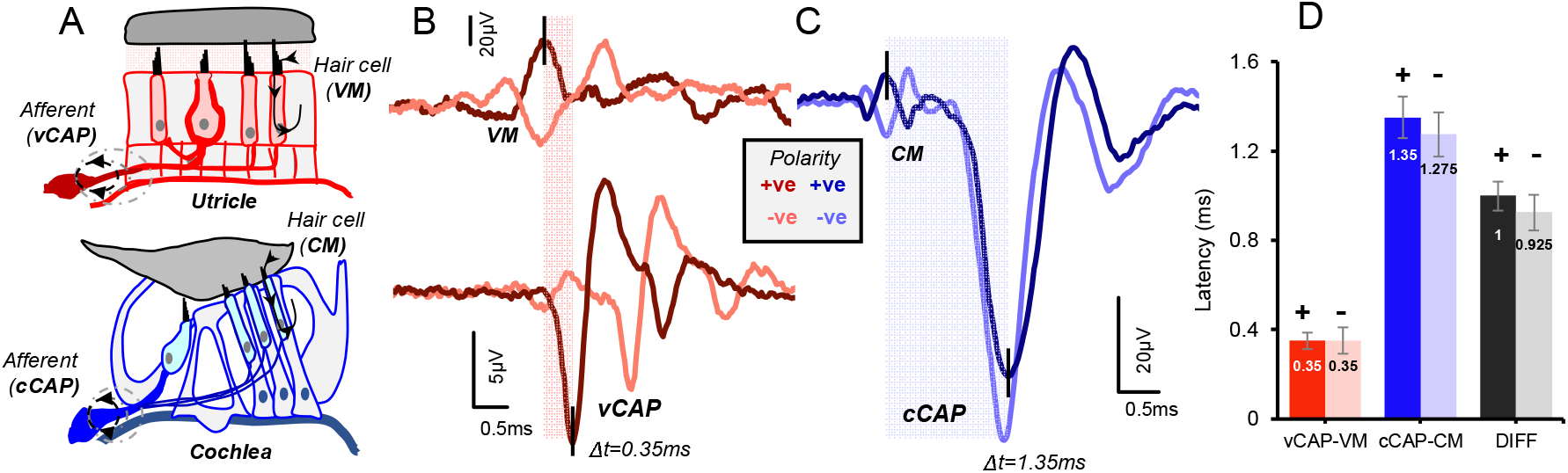
vCAP and cCAP response latencies. A) Schematic cross-section of the sensory regions of the utricle and the cochlea highlighting sensory generators from hair cells and afferent neurons. B) Simultaneous recordings of the VM and vCAP from a representative animal taken from the surface of the utricular macula and the facial nerve canal, respectively. The latency difference between the onset of the VM (first peak = P1) and the onset of the vCAP (first peak = N1) is 0.35 ms. C) Recordings of gross extracellular cochlear potentials from the same animal, in the opposite ear, with an early latency Cochlear Microphonic (CM) and a longer latency cCAP response, recorded from the round window. The latency difference between the CM (onset peak = P1) and cCAP (onset peak = N1) is 1.35ms. Note: both vestibular and cochlear responses in B) and C) were evoked by an identical 0.6ms Air-Conducted Sound (ACS) pulse, with a 0.3ms rise-fall time at a stimulus intensity 20dB above threshold. D) Latency comparisons across 5 animals. Absolute latency times between the vCAP and VM (red), the cCAP and CM (blue), and the difference between these two (DIFF) for each stimulus polarity; N=5.

### Forward Masking

We employed a forward masking paradigm to determine if the short latency vCAP could be reduced by depleting the RRP of synaptic vesicles in hair cells. Experiments followed the approach illustrated in Fig. 3A, where brief inter-aural acceleration stimuli were applied to the temporal bone (minishaker), and evoked vCAPs and VMs were recorded. vCAPs primarily reflect synchronized firing of many vestibular afferent fibres, while VMs primarily reflect mechano-electrical transduction (MET) currents at the electrode tip near the striola. The stimuli consisted of two pulses separated in time by the paired pulse interval (PPI) In the auditory nerve, which exhibits forward masking, the magnitude of the response is reduced as the PPI becomes small (Goldstein and Kiang, 1958). Example averaged vCAP and VM data are shown in Fig. 3B, using a 7ms PPI, where the first response was nearly identical to the second response. Shorter PPIs result in the second vCAP decreasing in amplitude. A plot of vCAP and VM amplitude vs PPI for this animal is shown in Fig. 3C, for PPIs ranging from 3.7ms to 107ms. The VM amplitude was completely unchanged by the PPI, analogous to results in the cochlea with the CM (Supplementary Figure S1). In contrast, the vCAP amplitude declined for PPIs <7ms, as would be expected as the second neural response falls within the relative refractory period of the action potential evoked by the first pulse.

**Figure 3:**
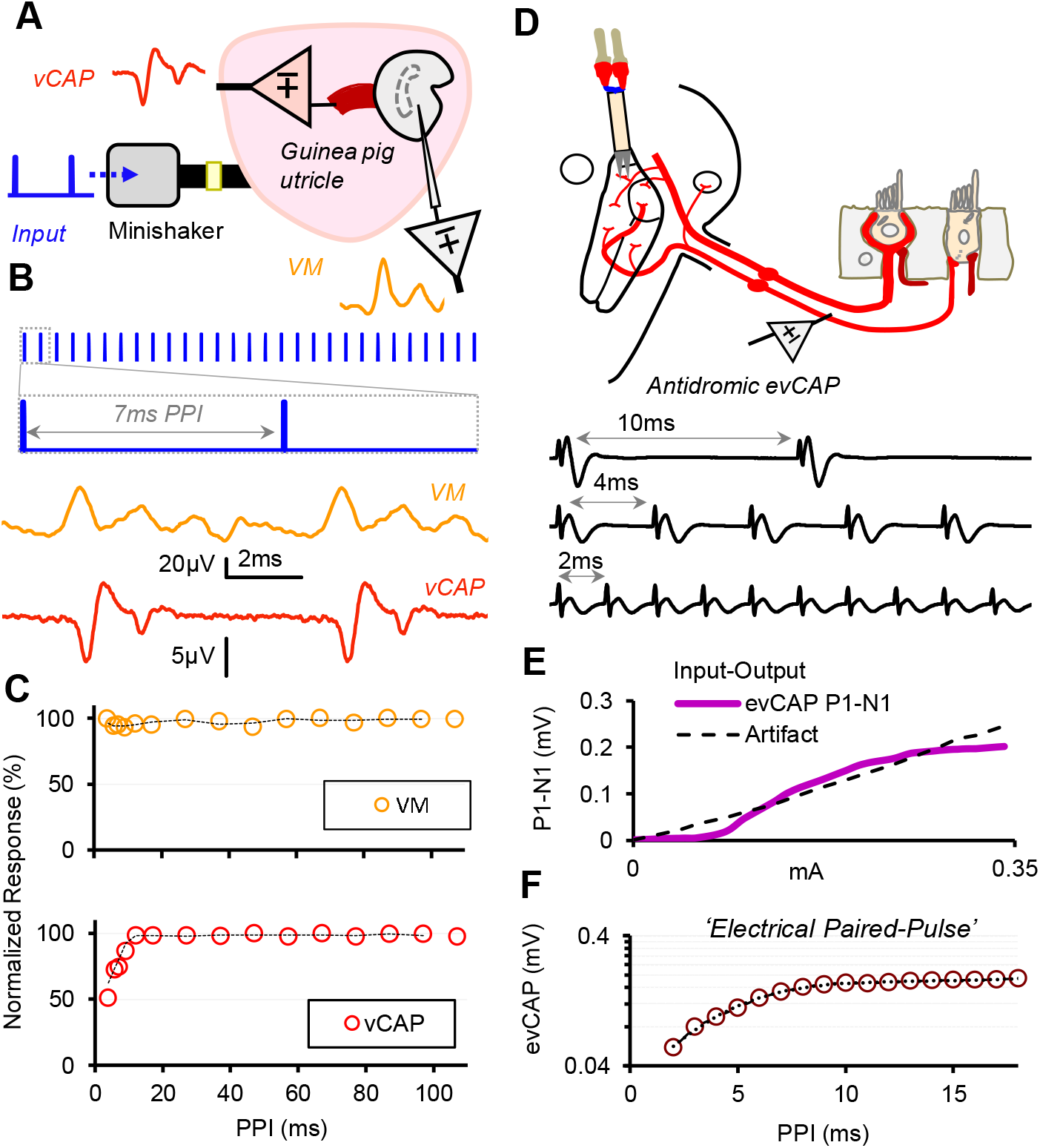
A) Bone-conducted vibration pulses were delivered to the guinea pig temporal bone via a minishaker, simultaneously evoking neural vestibular Compound Action Potentials (vCAPs) and transient hair cell potentials called Vestibular Microphonics (VMs). Vestibular PPIs were varied between 107ms and 3ms. B) Averaged VM and vCAP waveforms to 100 stimulus presentations for PPI stimuli of 7ms. C) VM and vCAP PPI response curves. vCAPs were insensitive to changes in PPI between 107ms and 10ms yet decreased with PPIs below ∼8ms. The VM amplitude was insensitive to changes across all PPIs D) Electrically-evoked vestibular Compound Action Potentials (evCAPs) were used to probe refractory properties of vestibular primary afferents. Bipolar current pulses were delivered to the proximal throughputs of the primary vestibular afferents at the central vestibular system, to evoke Antidromic evCAPs, recorded near the peripheral vestibular nerve branch. E) The Input-Output Function of the antidromic evCAP displays a saturating nonlinearity, whereas the early-latency electrical artifact is linear with increasing current intensities. F) To determine the relative refractory properties of the evCAP, the interval between successive 100μs biphasic current pulses or the electrical Paired-Pulse Interval (ePPI) was varied between 15ms and 2ms.

We further examined the refractory period of sensitive calyx bearing afferents using paired electrical stimulation. For this, electrically-evoked compound action potentials (evCAPs) were evoked using the paired pulse antidromic electrical stimuli (Ramekers et al., 2015) as illustrated in Fig. 3D. The antidromic evCAP response includes a saturating non-linearity as a function of increasing shock intensity, consistent with the sensitivity and Input-Output function of peripheral vestibular afferents to increasing mechanical and electrical stimuli (Pastras *et al*., 2018a). The relative refractory period of the primary calyx afferents was then characterised by varying the PPI of electrical shocks between 15 and 2ms (Fig. 3F) and monitoring the amplitude of the second evCAP response. Results demonstrated the second evCAP response amplitude exponentially declined with PPIs <8ms.

We compared vCAPs and cCAPs evoked by BCV paired-pulse stimuli to (Fig. 4) (∼15-20 dB above threshold)^1^ to determine if forward masking shown previously in auditory spiral ganglion neurons is also present in vestibular afferent neurons. The surgical approach is illustrated Fig. 4A, and example vCAP and cCAP signals are shown in 4B. BCV-evoked vestibular (vCAP, red), electrically evoked vestibular (evCAP, balck), and ACS-evoked cochlear (cCAP, blue) response amplitudes (N1-P1) are shown in Fig. 4D for paired pulse stimuli with PPIs varied from 3.7 to 107ms. Auditory cCAPs exhibited forward masking for PPIs <80 ms, while vestibular responses, whether evoked electrically or via BCV, only started to decline in magnitude only for PPIs <8-10ms, which is almost an order of magnitude less. This difference is highlighted in Fig 4E, where vCAP and cCAP responses to 8ms and 100ms PPI stimuli are presented (Fig. 4E).

**Figure 4:**
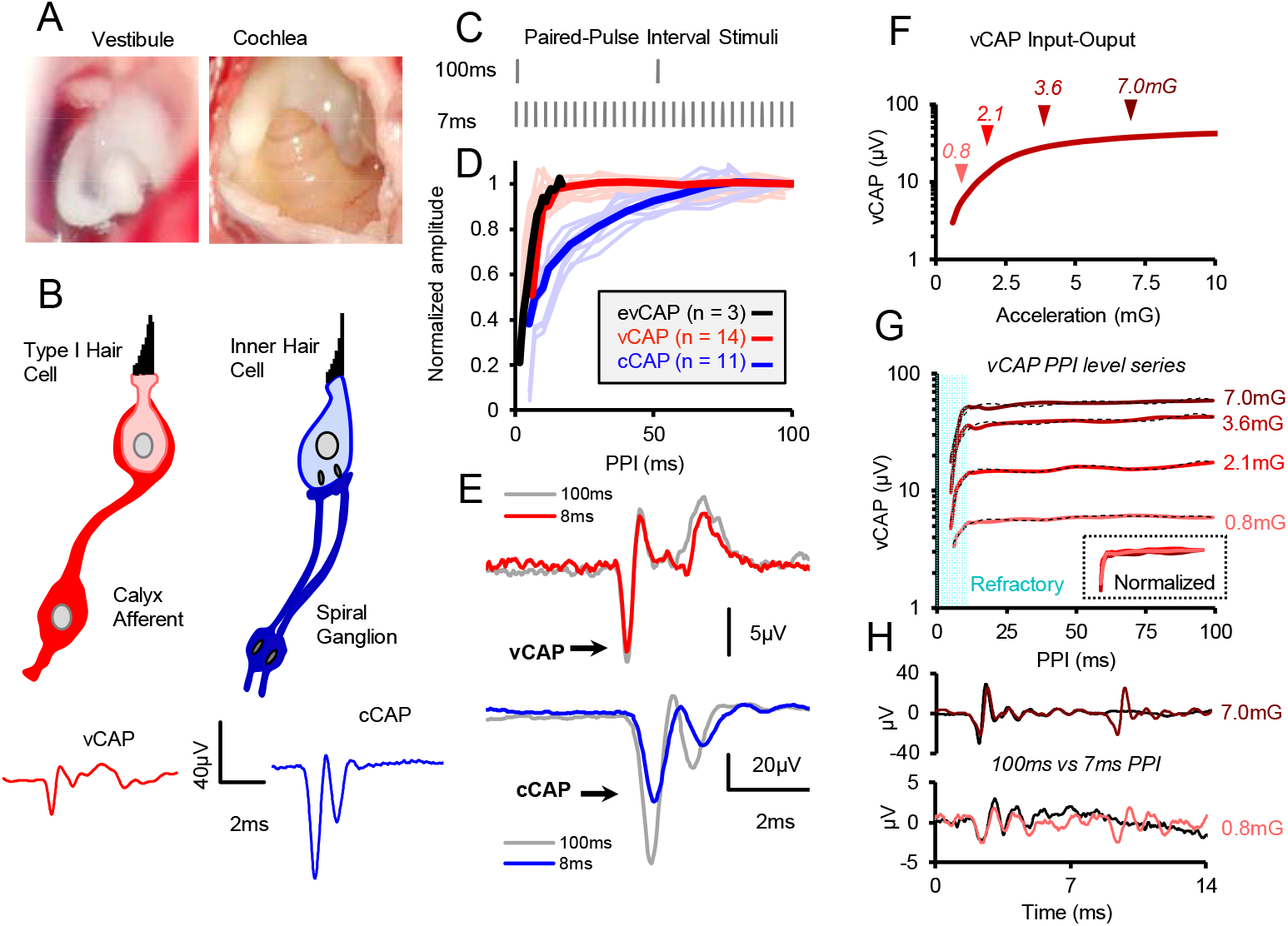
A) Photographs taken during surgery of the guinea pig utricle and cochlea in a representative animal. B) The mammalian vestibular system (utricle) contains Type I VHCs surrounded by unique chalice shaped calyx afferents, which synchronously fire action potentials to the onset of motion, producing vestibular nerve Compound Action Potential (vCAP) responses. The mammalian cochlea contains Inner Hair Cells which innervate multiple type-I SG Afferents, which evoke gross cochlear nerve Compound Action Potential (cCAP) responses to the onset of acoustic stimuli. C) Example Paired-pulse paradigms for 100ms and 7ms intervals between stimuli. D) Forward masking of vCAPs vs cCAPs. Normalized amplitude of mechanically evoked paired-pulse vCAPs (red) vs. mechanically evoked paired-pulse cCAPs (blue) vs. electrically-evoked vCAPs (black). E) Example records showing lack of forward masking in vCAPs and significant forward masking in cCAPs for a PPI of 8ms. F) vCAP Input-Output functions were measured to inform appropriate stimuli for G) vCAP PPI intensity curve testing. Mechanically evoked paired pulse vCAPs recorded at 4 different BCV stimulus levels, and vCAP normalized to max. amplitudes displayed in inset, revealed no forward masking associated with increased stimulus levels. H) Corresponding vCAP responses associated with 7.0mG and 0.8mG PPI intensity curves in G).

Finally, previous work in the cochlea has shown that auditory forward masking varies as a function of stimulus intensity, with greater forward masking occurring at higher stimulation levels (Harris and Dallos, 1979; Li et al., 2021), putatively due to increased depletion of the RRP (Eggermont, 2015). To investigate this effect, vCAP PPI response curves were recorded at 4 different BCV pulse stimulus intensities, corresponding to a broad section of the vCAP input-output growth function (Fig. 4F), just above threshold (0.8mG) to saturating intensities (7mG). Results reveal that stimulus intensity had no impact on vCAPs forward masking (Fig. 4G, H).

## Discussion

We present evidence that short latency vCAPs evoked by transient vibration in the guinea pig utricle arise from the ultrafast electrical component of NQ synaptic transmission at calyceal synapses. We begin with pharmacological evidence, followed with evidence based on timing, and finally with evidence based on forward temporal masking experiments.

It has been suggested that evolution of the vestibular calyx synapse in amniotes was driven by pressure to increase the speed of locomotor and reflex neural circuits, which depend heavily on the speed of inertial sensation by inner ear vestibular organs (Eatock, 2018). The present report examined the latency between a brief transient acceleration stimuli and action potentials evoked in sensitive calyx bearing afferent neurons in the guinea pig utricle. Results demonstrate the most sensitive vestibular afferents are remarkably fast, much faster than their auditory nerve counterparts. Here, we present evidence that the vestibular speed advantage arises from ultrafast NQ electrical synaptic transmission from Type I hair cells to their calyx partners (Contini *et al*., 2020). Direct electrical synaptic transmission improves speed for two major reasons. First, synaptic delay associated with chemical transmission is completely eliminated by electrical coupling, demonstrated here by the equivalence of electrically and mechanically evoked refractory periods (Fig. 4), and by the very short latency between MET currents and vCAP (VM vs. vCAP, Fig. 2). Second, electrical coupling does not suffer from depletion of presynaptic vesicles, evidenced here by the lack of forward masking in vCAPs (indistinguishable from the AP refractory period, Fig. 4). This was demonstrated by comparing paired-pulse mechanical stimuli to paired-pulse electrical stimulation of the nerve bundle, which distinguished pre- and post-synaptic mechanisms underlying the decline in vCAP magnitude for short PPIs. Remarkably, evCAPs recorded for paired electrical pulses were identical to vCAPs recorded for paired pulse mechanical stimuli, demonstrating that the reduction in vCAPs with short PPIs results from refractory properties of AP generation and not from hair cell synaptic transmission. Moreover, stimulus intensity had no effect on short latency vCAP forward masking, as would be expected for electrical NQ coupling at the calyx synapse. The fundamental innovation of the calyx was to eliminate chemical transmission in favour of electrical coupling that is ultrafast and immune to transmitter depletion (Fig. 4). The fact that auditory cCAPs were eliminated by the antagonist CNQX, while vCAPs were completely unchanged, offers additional strong evidence that glutamatergic quantal transmission is not required for short-latency vestibular responses (Fig 1). Importantly, hair cell microphonics were unaffected by CNQX suggesting that this blockade disrupted neurotransmitter release between hair cells and afferent neurons, and not mechanosensory hair cell function. This result alone, however, does not imply that neurotransmission at the calyx is non-quantal, because some other putative neurotransmitter, not affected by CNQX, may underlie transmission at the calyx. That said, when the lack of forward masking due to neurotransmitter depletion and latency, is coupled with the lack of CNQX effects, and the lack of evidence for any other major neurotransmitter present at this synapse, are taken together, evidence strongly argues against quantal transmission underpinning the vCAP response.

Short-latency responses are present only in calyx-bearing afferents contacting Type I hair cells in the striola (Curthoys *et al*., 2016). Three forms of synaptic transmission are present at these synapses: quantal, slow NQ K^+^ build up in the cleft, and ultrafast electrical NQ coupling. All three forms play a physiological role (Contini et al., 2022), but only the ultrafast electrical component is consistent with the present short-latency data. K^+^ build up is simply too slow (Govindaraju *et al*., 2023; Highstein *et al*., 2014; Lim *et al*., 2011) to account for the data. Quantal transmission is too slow, (Highstein et al., 1996; Li et al., 2014; Palmer and Russell, 1986), fatigable, vulnerable to forward masking (Harris and Dallos, 1979), and sensitive to CNQX. Present results rule out K^+^ build up and quantal transmission as candidate mechanisms but are entirely consistent with the ultrafast electrical NQ hypothesis.

Results raise the possibility that differences between auditory and vestibular responses to transient stimuli might offer a means to more clearly disambiguate auditory vs. vestibular evoked potentials. Transient stimuli delivered by BCV and ACS activate the otolith organs and the cochlea at the same time, leading to complex evoked potentials that reflect a mixture of auditory and vestibular responses. The auditory contribution to vestibular responses can be reduced by using acoustic masking (Jones and Lee, 2021), but our results suggest it might be possible to design even more discriminating stimuli (Fig. 2-4, & Supplementary Figure 3).

## Acknowledgments

This work was supported by a Macquarie University Research Fellowship, MQRF0001126 (CJP), and a NIH DC 006685 (RDR).

## Footnotes

1 Note, PPI response curves in both the cochlea and vestibular system were identical for both BCV (bone-conducted vibration) or ACS (air-conducted sound) pulses (Supplementary Fig. S2).

