## Supplementary Information for "Evidence that ultrafast non-quantal transmission underlies short-latency vestibular evoked potentials"

**Supplementary Note 1.** Given forward masking in the cochlea is largely dependent on glutamate neurotransmitter depletion from the inner hair cell, there should be minimal effect on the level of transduction currents entering the hair cells and the gross extracellular receptor potential. This was tested in our experimental setup, with direct comparisons of the air-conducted sound (ACS) Cochlear Microphonic (CM) to the cochlear nerve Compound Action Potential (CAP) during Paired-pulse interval (PPI) stimuli ranging from 160-2ms. Results reveal that the auditory neural CAP, but not the auditory hair cell CM is sensitive to forward masking.

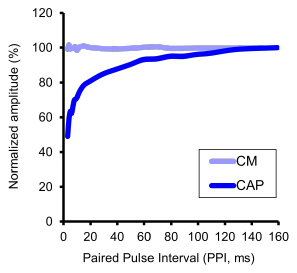

**Supplementary Figure 1 (S1).** Comparisons of mechanical PPI effects on the Cochlear Microphonic (CM, light blue) vs. the cochlear nerve Compound Action Potential (CAP, blue). Measurements taken from the round window of the anaesthetized guinea pig.

**Supplementary Note 2.** Mechanical PPI responses were generally recorded using ACS for the cochlea, and bone-conducted vibration (BCV) for the vestibular system. Given, the differences in stimuli, such as complex 3D transmission modes through the skull for BCV, attempts were made to separate any stimulus related effects by comparing PPI response curves for both sound and vibration in the cochlea and vestibular system (utricle). Results from a representative animal reveal no significant differences between PPI effects based on stimulus in the cochlear and vestibular compound nerve response using BCV or ACS at 20dB above threshold.

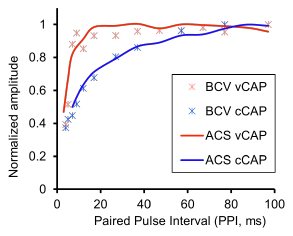

**Supplementary Figure 2 (S2).** The mechanical PPI effects on vestibular and cochlear CAPs do not change across BCV and ACS pulsatile stimuli. vCAP measurements were taken from the facial nerve canal, and cCAP responses were recorded from the round window in the same ear of the anaesthetized guinea pig.

**Supplementary Note 3.** Given the translational appeal of using differences in PPI response curves to separate vestibular from cochlear responses, we wanted to determine if such effects were reproducible with changes in recording location. Results reveal that the vestibular PPI response curve is consistent when recorded from near-field locations, such as the facial nerve canal (FNC), or from far-field ‘clinical’ locations, such as the vertex.

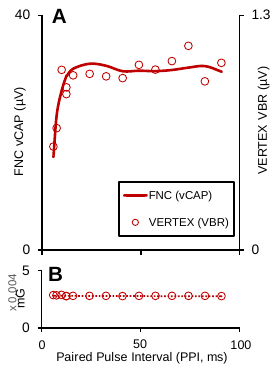

**Supplementary Figure 3 (S3).** Vestibular nerve mechanical PPI effects are consistent across near- (invasive) and far-field (non-invasive) recording locations. a. The facial nerve canal (FNC) vestibular Compound Action Potential (vCAP) and vertex vestibular brainstem response (VBR) have identical PPI response curves to b. pulsatile BCV acceleration transients associated with PPIs between 109-7ms.
